# A phylum-wide survey reveals multiple independent gains of head regeneration ability in Nemertea

**DOI:** 10.1101/439497

**Authors:** Eduardo E. Zattara, Fernando A. Fernández-Álvarez, Terra C. Hiebert, Alexandra E. Bely, Jon L. Norenburg

## Abstract

Animals vary widely in their ability to regenerate, suggesting that regenerative abilities have a rich evolutionary history. However, our understanding of this history remains limited because regeneration ability has only been evaluated in a tiny fraction of species. Available comparative regeneration studies have identified losses of regenerative ability, yet clear documentation of gains is lacking. We surveyed regenerative ability in 34 species spanning the phylum Nemertea, assessing the ability to regenerate heads and tails either through our own experiments or from literature reports. Our sampling included representatives of the 10 most diverse families and all three orders comprising this phylum. We generated a phylogenetic framework using sequence data to reconstruct the evolutionary history of head and tail regeneration ability across the phylum and found that while all evaluated species can remake a posterior end, surprisingly few could regenerate a complete head. Our analysis reconstructs a nemertean ancestor unable to regenerate a head and indicates at least four separate lineages have independently gained head regeneration ability, one such gains reconstructed as taking place within the last 10-15 mya. Our study highlights nemerteans as a valuable group for studying evolution of regeneration and identifying mechanisms associated with repeated gains of regenerative ability.

## Introduction

Regeneration, the ability of an organism to regrow a body part following traumatic loss, is a fascinating phenomenon that occurs in many animal groups. Regeneration of specific body structures (e.g., heads, tails, appendages) and regeneration from a tiny fragment (whole body regeneration) are both found scattered across metazoan phylogeny [1–3]. Lineages that are sister to the Bilateria (e.g., Porifera, Ctenophora and Cnidaria), generally possess high regenerative ability, suggesting that early animals likely also had high regenerative ability [1,4,5]. Within the Bilateria, however, regenerative ability is extremely variable, indicating a complex pattern of regeneration evolution. Within phyla of Ecdysozoa, regenerative abilities are generally very restricted, with limb regeneration in Arthropoda being the main exception [6]. Both across and within most other bilaterian phyla, however, regeneration ability ranges widely. Species with extensive regenerative abilities are common in the Xenacoelomorpha [7,8], deuterostome phyla such as Echinodermata [9], Hemichordata [9], and Chordata [10,11], and spiralian phyla such as Platyhelminthes, Mollusca, Annelida and Nemertea [12]. However, most of these same phyla also include representatives with modest or even extremely limited regenerate ability. Thus, even a cursory overview of regeneration ability indicates that the pattern of regeneration evolution is complicated.

Determining where increases and decreases in regenerative ability have occurred across animals is important for understanding how regeneration evolves. Doing so reveals the pattern of regeneration evolution and allows for identifying developmental processes potentially responsible for changes in regenerative ability. Currently, knowledge of regeneration ability remains extremely sparse for most animal phyla, limiting understanding of the pattern of regeneration evolution. Furthermore, while a considerable amount of basic and applied research has focused on the developmental mechanisms involved in regeneration [3,5,13–16], the vast majority of this research has focused on a small set of species that are deeply diverged from one another, limiting understanding of the mechanisms of regeneration evolution. Only a few studies have compared regeneration between closely-related species that vary naturally in their ability to regenerate homologous body parts. Such studies have proved extremely informative, demonstrating for example that variation in regenerative ability can result from just a few changes in key molecular and developmental processes [17–21]. Expanding the number of groups in which regeneration increases (i.e., gains) and regeneration decreases (i.e., losses) are well documented is likely to provide new insights into regeneration evolution.

Losses of regeneration have been inferred in several animal groups [22]. As mentioned above, it is likely that early animals could regenerate well, and that restrictions in regeneration ability evolved later within Bilateria. Comparisons of regenerative ability between phyla are difficult to interpret, however, due to issues regarding homology (how can regenerative ability across species be compared if the structures being regenerated have unclear homologies? [3]) and because there may be considerable variation within each of the phyla being compared (such that the ancestral states for the phyla being compared are unclear). To date, only one study has analyzed regenerative ability across an entire phylum to reconstruct ancestral states and formally identify putative gains and losses [23]. This recent study on Annelida inferred the ancestral state for the phylum as having both anterior and posterior regenerative ability and also identified many losses, of both anterior and posterior regeneration ability. Despite the large dataset of several hundred annelid species, this study identified no gains of regeneration.

Evidence for clear increases of regenerative ability is thus far very limited. Limb regeneration in arthropods likely represents a gain of regenerative ability, given the extremely limited regenerative abilities of most ecdysozoans [5]. Limb regeneration in salamanders and tail regeneration in lizards may also each represent gains, given the weak or absent regeneration of these same structures in the closest relatives of these groups [24]. Although these putative gains are interesting, they would have happened at phylogenetic nodes so deep that comparative approaches have little chance to uncover meaningful mechanistic insights into their underlying causes. In contrast, identifying more recent gains of regenerative ability would potentiate studies of the proximate (developmental) and ultimate (evolutionary) causes behind regeneration enhancements. To date, no comparative studies have yet uncovered clear gains of regeneration across relatively close relatives.

Ribbon worms (phylum Nemertea) are a promising group for investigating the evolution of regeneration. Nemerteans are elongated, primarily marine predatory worms with highly flexible bodies. The phylum has a reputation for possessing high regenerative ability that is based almost entirely on the remarkable regeneration abilities of one species: *Lineus sanguineus* Rathke. This species is unquestionably one of the champions of animal regeneration, possessing some of the highest regenerative abilities known among animals [12,25]. Individuals of this species can be repeatedly amputated until the resulting worms that regenerate are just 1/200,000^th^ of the volume of the original individual. A complete animal can regenerate not only from a thin transverse slice of the body, but from even just one *quadrant* of a thin slice (with a large majority of the fragment’s surface area being wound surface)[26]. Although the regenerative abilities of this species are spectacular and well-described, they do not appear to be typical for this phylum. Nemertea comprises ^~^1200 species and regenerative ability has been described from a few additional species, yet none comes close to the remarkable abilities of *L. sanguineus*. Furthermore, there has likely been a publication bias against reporting findings from poor regenerators, as is suggested for other groups [22,23]. Thus, regenerative ability appears to be variable among nemerteans, but the phylogenetic pattern within this phylum remains very poorly understood.

When placed in the context of current understanding of nemertean phylogeny, the limited regeneration data available for nemerteans yields at best a blurry picture of regeneration evolution in this phylum[12]. Nemerteans have traditionally been placed into three orders: Palaeonemertea, Hoplonemertea, and Pilidiodophora (Heteronemertea). Palaeonemertea are likely a paraphyletic assemblage of basal lineages [27]. No regeneration data is available in the literature for any species in this order. Hoplonemertea is a well-supported clade, with most species reported in the literature to have quite limited regenerative abilities; unfortunately, most reports of regeneration are presented as blanket statements, without specifying the species that have been examined [28]. Pilidiophora (Heteronemertea) is a large and well-supported clade [27,29–31], and many species are frequently cited as examples of nemerteans with outstanding regenerative ability [25,32,33]. However, all of these “many species” [26,28,34–40] have now been synonymized to *Lineus sanguineus* [41–43]. Thus, regeneration data remain very cursory across Nemertea but do suggest that high regenerative ability – in particular, the ability to regenerate a head – may be uncommon in the phylum. Systematic testing of regeneration ability of well identified species is clearly needed to resolve the pattern of regeneration evolution in this phylum.

In this study, we addressed the question: what is the broad pattern of regeneration evolution within the Nemertea? We conducted a survey of regenerative abilities among species from across the phylum, performing new regeneration experiments on 22 species and obtaining information from the literature for 12 additional species. Using nucleotide sequence data we collected ourselves or obtained from public databases, we generated a phylogenetic framework and mapped the results of this regeneration survey. We estimated ancestral states for all nodes in our phylogeny and reconstructed the pattern of gains and losses of regenerative ability across the phylum.

## Materials and Methods

### Regeneration survey

Nemerteans were collected worldwide between the years 2012 and 2014, including along the Atlantic and Pacific coasts of the United States, along the Atlantic coast of Argentina, along the Atlantic coast of Spain, and along the coasts of New Zealand’s South Island. Tables S1-S2 in the electronic supplementary material provide a full list of location, collectors and taxonomic nomenclature. Due to the patchy distribution and abundance of species, the sampling was opportunistic, but we aimed to collect all major lineages within the phylum.

For regeneration experiments, we bisected worms by cutting transversely, generating an anterior and a posterior fragment. We cut at two different possible transverse planes, cutting at ^~^1/3 the total body length (all species) or cutting at ^~^2/3 the total body length (in species in which several specimens were available). In all species, amputation planes were posterior to the mouth and the cephalic nervous system (brain and cerebral organs). Samples sizes ranged from 1 to >40 cut animals per species (resulting in twice this many fragments). We maintained amputated specimens in seawater, without food, and scored survival and externally visible post-amputation phenotypes. We used a series of standard morphological and behavioral criteria (detailed in the electronic supplementary material) to determine whether amputated specimens showed evidence of posterior and/or anterior regeneration of the missing end, as well as the time to complete regeneration (when present). Regeneration of each type was scored as present even if not all experimental individuals completed all landmarks. When multiple individuals were scored, approximate times for each landmark were summarized as a range, except for completion of regeneration, where the fastest cases were reported, and survival without regeneration, where the longest survival times were reported. Experimental specimens showing clear signs of poor health or abnormal development were excluded from timing estimations.

We expanded the number of nemertean species in our regeneration dataset using literature searches as described in [23]. Data were included in our dataset only if regeneration results were unambiguous, based on amputations similar to those from our own experiments, and involved identifiable, valid species.

### Molecular marker sequencing

DNA was extracted using a DNeasy 96 Blood & Tissue Kit (69581, Qiagen) from at least one individual of each species used in regeneration experiments. Whenever possible, the extraction was made from individuals that had undergone the amputation experiments; when that was not possible, we used conspecific individuals from the same field collection. We amplified by PCR fragments of four genes, cytochrome oxidase subunit I (COI), 16S ribosomal RNA (16S), small subunit ribosomal RNA (18S) and large subunit ribosomal RNA (28S). Primers sequences and PCR parameters are detailed in the electronic supplementary material. PCR products were purified using ExoSAP-IT (Thermo-Fisher), and sequenced in paired reactions using respective forward and reverse primers with the BigDye™ Terminator v 3.0 Cycle Sequencing Kit ver. 3.0 (Applied Biosystems). Sequencing reaction products were analyzed using an ABI Prism 3730xl Genetic Analyzer capillary sequencer (Applied Biosystems). For several species of *Lineus*, sequences were obtained from published transcriptomes [43]. In the few cases in which we had regeneration data (either from our experiments or the literature) but no associated sequence data, we retrieved relevant sequence data available from NCBI. All sequence data were deposited at NCBI (see Table S3 of the electronic supplementary material).

### Sequence alignment and phylogenetic reconstruction

Sequence quality assessment, assembly, and alignment and phylogenetic reconstruction were performed using the Geneious 8.1.9 platform [44]. Sequences were aligned into a multiple sequence alignment (MSA) for each marker using the MAFFT algorithm [45], curated by eye and concatenated. The concatenated MSA was used as input for RAxML v 8.2.11 [46], set up to perform 100 rapid bootstrap inferences followed by a thorough maximum likelihood search, using a General Time Reversible (GTR) model with gamma-distributed rate heterogeneity. The MSA was divided into six partitions, each run with different models: three partitions were used for the protein coding marker COI (one partition for each codon position) and one partition was used for each of the rRNA markers (16S, 18S and 28S). The inference was run first without topological constraints (“unconstrained”), and then re-run with alternative topological constraints reflecting different hypotheses about deep phylogenetic relationship within the Nemertea (see the electronic supplementary material for details). We also performed Bayesian inference from the MSA using MrBayes 3.2.6 [47], specifying a GTR model with 4 categories of gamma distributed rate heterogeneity and a proportion of invariant sites, and no topological constraints. Four heated chains were run for 1,100,000 steps and subsampled every 200 steps; the initial 100,000 steps were discarded as burn-in.

### Ancestral trait estimation by maximum likelihood

Best scoring trees from each analysis were used as phylogenetic frameworks for character mapping and ancestral trait estimation. We coded regeneration ability as two binary variables, presence/absence of anterior regeneration and presence/absence of posterior regeneration, and generated a matrix that included these two variables for each taxon (species or population) in our MSA. We used the *ace* function from the *ape* package [48], which models discrete trait state evolution as a Markovian process [49]. This function incorporates phylogenetic tree branch length information to estimate the rates of change of the trait and the likelihood for each character state at every node of the tree, including the basal node [50]. A two-parameter model was specified allowing for separate calculation of the rate of gain (0→1) and rate of loss (1→0). We repeated this procedure for all the trees inferred using the different constraint sets (see above). All analyses were run within the R computing environment [51].

## Results

### Regeneration survey

We collected and performed regeneration experiments on 22 nemertean species: 4 species of Palaeonemertea, 6 species of Hoplonemertea and 12 species of Pilidiophora. We also obtained data from the literature for 13 additional species, producing a final regeneration dataset of 35 species (see Table S4 of the electronic supplementary material). Although the number of species in our dataset is a small fraction of the known nemertean diversity, it nonetheless represents 10 of the most diverse families, and spans all three orders (2 out of 3 paleonemertean families, 4 out of 20 hoplonemertean families and all 4 pilidiophoran families [52]).

Outcomes of regeneration experiments for each species are described in the electronic supplementary material. Overall, we found that in all species, most individuals (>90%) survived the initial amputation, and fragments usually healed the wounds within 5 days post-amputation (dpa; Table S4). All species were able to complete posterior regeneration (Figure 1). However, most species (27/35) were not capable of regenerating a complete head (including a brain), despite many species being able to survive without the missing structures for several weeks or months (Figure 1, Table S4). Location of the amputation plane (at either 1/3 or 2/3 of the body length) had no influence on regeneration success for those species in which cuts were made at both planes.

**Figure 1:**
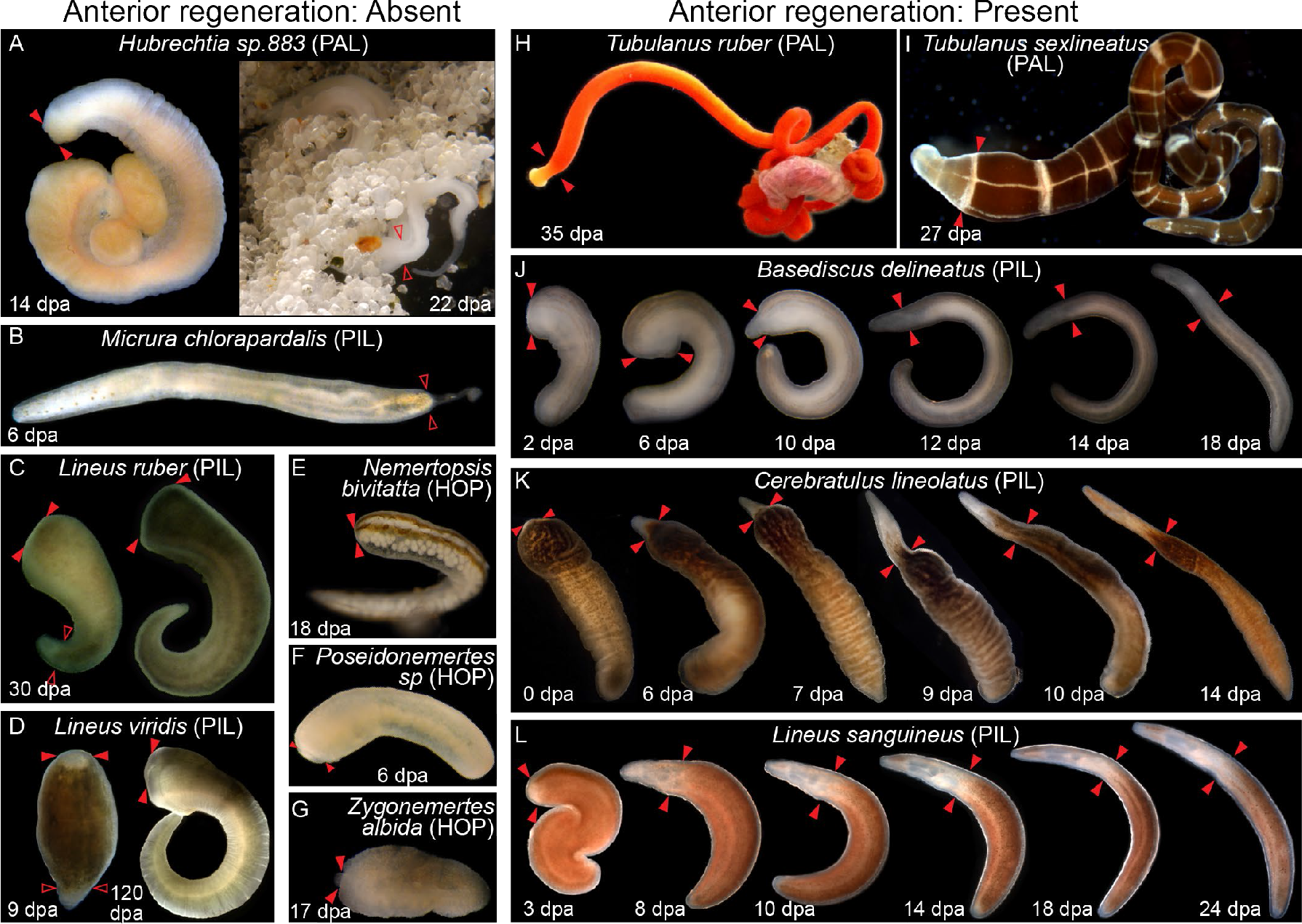
Photodocumentation of regenerative ability in representatives of Nemertea. Individuals shown were amputated posteriorly and/or amputated anteriorly (time since amputation is shown at bottom). Posterior regeneration ability was detected in all nemertean species in our study; anterior regeneration was detected in only eight of these species. Shown here are photos of representative individuals undergoing successful posterior regeneration (A-D, J), failed anterior regeneration (A-G) and successful anterior regeneration (H-L). Species in which anterior regeneration was scored as absent are on the left; species in which anterior regeneration was scored as present are on the right. Plane of posterior amputation is indicated by paired, empty arrowheads (A-D, J); plane of anterior amputation is indicated with paired, filled arrowheads (A, C-L). Panels J-L show regeneration time series of the same experimental individual over time. All individuals within the same panel are at the same scale. Anterior is left, upper left, or up. PAL: Palaeonemertea; HOP: Hoplonemertea; PIL: Pilidiophora; dpa: days post-amputation.

Successful head regeneration was found in four species where it was previously unreported: *Tubulanus ruber* and *T. sexlineatus* (Paleonemertea), *Baseodiscus delineatus* and *Cerebratulus lineolatus* (Pilidiophora). We also observed head regeneration on *Lineus sanguineus* (Pilidiophora) from several locations. Our literature review added *Lineus pseudolacteus* [53] and *L. pictifrons* [35] (Pilidiophora), and *Prostoma graecense* [54] (Hoplonemertea) to this list.

### Sequencing and phylogenetic framework inference

We collected sequence data for four phylogenetic markers (COI, 16S, 18S and 28S) for species in our regeneration dataset, collecting some novel sequence data and obtaining additional sequence data from available databases. Our sequence dataset comprised 114 new Sanger sequences, 55 new RNAseq-based sequences, and 35 sequences retrieved from the NCBI nr/nt database. New sequences have been deposited at NCBI (see Table S5 of the electronic supplementary material for accessions).

Using automated alignment followed by manual curation and trimming, we generated a multiple sequence alignment that was 8123 bp long and had 3921 distinct alignment patterns (unique columns). We inferred phylogenetic trees using maximum likelihood searches (RAxML trees) and Bayesian inference (MrBayes tree). When no topology constraint was enforced, both methods found mostly congruent trees, with monophyletic Palaeonemertea, Hoplonemertea and Heteronemertea (Figure 2 and Figures S1-S6 in the electronic supplementary material). The only difference between the inferences was that the RAxML tree grouped Palaeonemertea and Heteronemertea into a sister group to Hoplonemertea, while in the MrBayes tree the branching order of the three clades was not resolved. When topology constraints were enforced (see Methods), the resulting inferences differed only in the enforced bipartitions, but the internal topology of the remaining clades did not differ from the unconstrained trees. Our results are similar overall to those of previous studies [29–31,55,56], and are further described in the electronic supplementary material.

**Figure 2:**
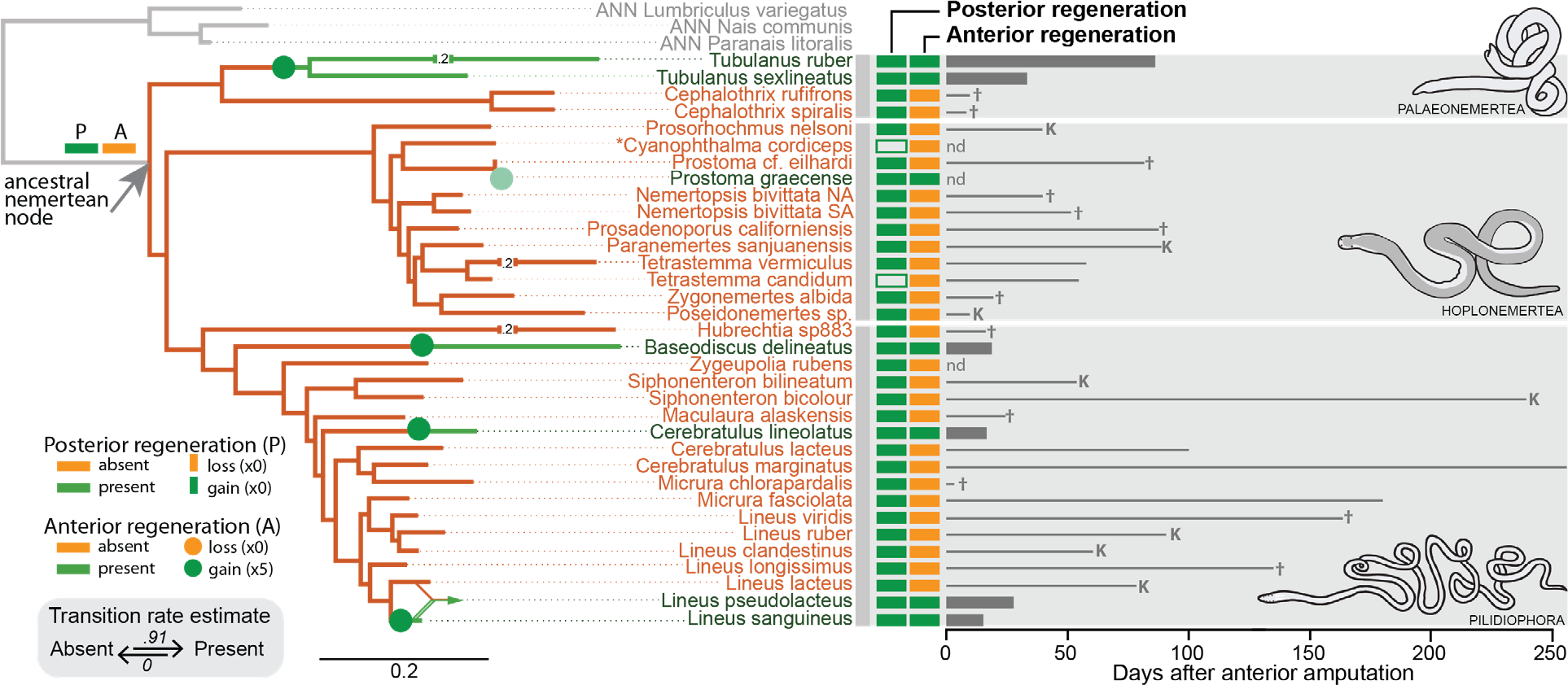
Phylogenetic distribution of regenerative abilities in the phylum Nemertea. Maximum likelihood molecular dendrogram inferred from a multiple sequence alignment of two mitochondrial (COI, 16S) and two nuclear (18S, 28S) markers, analyzed on RAxML with a GTRGAMMA partitioned model and constrained to the Pilidiophora hypothesis (see Methods). Branch colors represent the estimated state for anterior regeneration ability (orange: absent, green: present); grey branches lead to three outgroup species (ANN: annelids). Numbers on broken branches indicate abridged distances. Green circles represent evolutionary transitions; the fifth, lighter green circle indicating a gain in *Prostoma graecense* is placed based on the position of *P. eilhardii* in our analyses. The location of the transition along a given branch is arbitrary. The converging lines leading to *Lineus pseudolacteus* represent the possible hybrid origin of this species. Orange and green boxes by species names show regenerative abilities (orange: absent, green: present; outlined boxes represents putative results) scored experimentally or taken from the literature. Bars to the right indicate either days to complete anterior regeneration. Lines to the right indicate longest survival time in days without a head, in species where absent. For species experimentally assessed, † indicates natural death while K indicates sacrifice or accidentally death of the longest surviving individual. Data from literature reports with no survival time are marked “nd”. *Lineus pictifrons* is not included in this diagram because there are no available data about the phylogenetic position of this species.

### Ancestral character estimation analysis

Given that posterior regenerative ability was invariant (present throughout our dataset), no further formal analyses were performed for this trait. As for anterior regenerative ability, we found that analyses based on any of the inferred phylogenetic trees gave the same qualitative results. Specifically, all analyses strongly support the absence of anterior regeneration ability at the root node of Nemertea (Figure 2). Runs on alternative topologies yielded only minor differences in the resulting log-likelihoods (ranging from −12.60 to −12.54) and transition rate parameters (ranging from 0.9 to 1.4 for gains, and 0 for losses).

Based on the likelihood of anterior regeneration being present or absent at each node of the trees, our analyses suggest at least four independent gains of anterior regeneration across the phylum (Figure 2): one in the *Tubulanus* lineage, one in the *Baseodiscus delineatus* lineage, one in the *Cerebratulus lineolatus* lineage and one in the *Lineus sanguineus* lineage.

## Discussion

A fundamental step towards understanding how and why regeneration abilities evolve is to estimate where and when changes in regenerative abilities have occurred across the animal phylogeny. Currently, however, sparse sampling and deep phylogenetic distances between most species being compared has severely limited our ability to infer patterns and mechanisms of regeneration evolution. To date, only a tiny fraction of animal species has been assessed for regenerative ability and, of the approximately 35 animal phyla, rigorous evolutionary reconstructions of regenerative ability have previously been performed across only a single phylum, Annelida [23]. To expand our knowledge of the evolution of regeneration abilities across animals, we performed a phylum-wide study of regenerative ability in the phylum Nemertea, the ribbon worms. We gathered regeneration data and sequence data from 35 species spanning the phylum and reconstructed the pattern of regeneration evolution across the phylum.

We found that most nemertean species assessed for regeneration ability are unable to regenerate a complete anterior end (i.e. a head that includes a brain) and we reconstruct that the last common ancestor of nemerteans likely lacked anterior regeneration ability. Thus, although Nemertea includes a few species with outstanding regeneration capabilities, including whole body regeneration, and although posterior regeneration is widespread, our analyses suggest that anterior regeneration ability is a derived feature in this phylum.

Our evolutionary analyses indicate that the ability to regenerate a head evolved independently at least four times within Nemertea. These gains represent some of the most clearly documented increases of regenerative ability known among animals and are the first well-documented gains of head regeneration ability among animals. One of these gains of anterior regeneration, involving *Lineus lacteus, L. pseudolacteus* and *L.sanguineus,* appears to be evolutionarily very recent [43], making these species an excellent system in which to further investigate regeneration evolution. Our findings contrast strongly with the pattern of regeneration evolution inferred in Annelida, another group of worms that are relatively closely related to Nemertean, and the one other group in which evolution of regeneration has been inferred at a phylum-wide scale [23]. Thus, our study highlights that evolutionary histories of regeneration may differ markedly across phyla.

### All nemertean species investigated can reform a posterior end, but most cannot regenerate a complete anterior end

We found that all species investigated were able to reform a posterior end but that most (27/35) species were not able to regenerate a complete head (including a brain), even if individuals survived several weeks or months after amputation. This general pattern was previously suggested by several nemertean researchers based on more limited and largely unpublished observations [28,36,52,57]. Our study, which includes far more species than previously considered and broader coverage across the phylum, supports these early inferences and provides strong evidence that anterior regeneration ability is uncommon in Nemertea.

That the ability to reform the posterior end is prevalent in nemerteans is not unexpected. Posterior regeneration appears to be widespread, and far more common than anterior regeneration, in many animal groups, including annelids, platyhelminths, mollusks, and vertebrates [1,12]. Several hypotheses could explain the higher prevalence of posterior regeneration as compared to anterior regeneration, such as different selective forces acting on the replacement of anterior and posterior tissues, or high pleiotropy between posterior regeneration and growth [1]. Although we found evidence of reformation of the posterior end for all species in our dataset, it should be noted that scoring for the reformation of the posterior end is challenging in nemerteans, especially when assessments are limited to external observation (as was the case in our study). Many nemertean species lack any morphologically distinctive posterior structures, and, in the absence of these, the only scorable posterior traits are the anus and a diffuse posterior growth zone [58]. While these features can be inferred based on morphology, observing defecation and/or elongation of the newly formed posterior end is the only definitive way to determine that posterior regeneration is indeed complete. Unfortunately, many species will not feed in laboratory settings, and thus do not defecate or grow noticeably in such a setting. In our study we scored for the reformation of the posterior end based on the reappearance of any distinctive posterior structures (if these were present in the species), of the anus, and of the diffuse posterior growth zone. However, the possibility remains that posterior regenerative abilities have been overestimated in our survey and thus future studies involving feeding (so that defecation and posterior elongation can be scored) and histological analysis (to definitively score for anus formation) should be performed to confirm our results.

The ability to regenerate anteriorly was found to be far more limited across Nemertea than the ability to regenerate posteriorly. Eight of 35 species assessed for regeneration were found capable of anterior regeneration. Of these, four were previously known; this study represents the first report of anterior regeneration ability in four additional species. The anteriorly-regenerating species *Lineus sanguineus* (including forms described as *L. nigricans, L. socialis, L. vegetus* and *L. bonaerensis*) plus the hybrid species *L. pseudolacteus* [43] were previously described as possessing outstanding regenerative abilities. We also found reports in the literature of complete head regeneration after amputation, at a narrow range of positions, for the hoplonemertean *Prostoma graecense* [54] and the pilidiophoran *Lineus pictifrons* [35], but were unable to collect specimens for experimental verification. Our work is the first to report of the presence of anterior regenerative ability in the palaeonemerteans *Tubulanus ruber* and *T. sexlineatus*, and the pilidiophorans *Baseodiscus delineatus* and *Cerebratulus lineolatus*.

Our confidence in accurately scoring species for anterior regeneration ability is high for several reasons. First, unlike posterior regeneration, anterior regeneration in nemerteans involves clearly recognizable intermediate stages, including the formation of a blastema that is evident morphologically (being composed of a tightly packed mass of cells with low pigmentation). Second, amputation of the head removes the mouth, and thus halts the ability to feed, such that food availability cannot influence regeneration output. Therefore, we expect high accuracy in detecting both the presence and the absence of anterior regeneration ability. This being said, evidence for the absence of anterior regeneration is necessarily weaker than evidence for the presence of anterior regeneration, especially in those species for which only a few specimens were available for experimental assessment. Thus, we hope that future regeneration studies will be performed on a broad range of nemerteans to corroborate and expand on our findings.

### The nemertean last common ancestor likely lacked anterior regeneration ability

Reconstructing the regenerative abilities of the ancestor of animal phyla is a key step towards understanding the broad pattern of regeneration evolution in animals. Knowing the ancestral regenerative condition is necessary to polarize changes in regeneration ability within a phylum (e.g., to determine whether regeneration gains or regeneration losses have occurred) and is critical for making meaningful comparisons of regeneration between phyla. We thus were interested to use our data to address the question: was the ancestral nemertean capable of regenerating a complete head? We found anterior regeneration ability to be absent from most species tested across the phylum. However, among the Palaeonemertea (representing the basal-most lineages of Nemertea), two of the four species assessed (the two *Tubulanus* species) were found to be capable of anterior regeneration, such that it was not obvious what the ancestral regeneration ability might have been. We therefore performed a formal analysis to estimate the anterior regeneration ability state at the base of Nemertea.

Our ancestral character estimation analyses consistently yielded a zero likelihood for anterior regeneration being present in the last common ancestor of Nemertea. This outcome was found even when considering alternative topologies (including one where tubulanids represent the most basally branching lineage). This result of anterior regeneration being absent at the base of Nemertea stands in sharp contrast to the widespread regenerative capabilities of basal bilaterians [1,5] and also to results from a similar analysis made on the phylum Annelida that found strong support for anterior regeneration being present at its basal node [23]. The contrast between Nemertea and Annelida is all the more striking as both phyla are within the same bilaterian subclade, Spiralia, and both are soft-bodied elongated animals (“worms”) with a similar level of body complexity. The closest relatives of Nemertea have been relatively poorly sampled for regenerative ability, but regeneration of particular body regions is known from the three most closely related phyla. Based on recent phylogenetic studies, Nemertea is sister to Phoronida, a few of which can regenerate the primary body axis [59], and Brachiopoda, some of which can regenerate the shell, lophophore and pedicle [60]. These three phyla together form a clade sister to Mollusca [61], some of which can regenerate the foot, tentacles, mantle, and eyestalks but which, as a group, does not appear to have widespread, extensive regenerative abilities [12]. Also within Spiralia are the Platyhelminthes, with both highly regenerative representatives and weakly regenerating representatives. Even though more extensive regeneration surveys and formal ancestral state estimation are needed for these other spiralian phyla, placing our results for nemerteans in the broader context of our current knowledge suggests that the Spiralia subclade of bilaterian animals has had a rich evolutionary history with respect to regeneration and that regenerative ability was highly variable even at deep nodes within this clade.

### Head regeneration ability evolved independently at least four times within Nemertea

The most unexpected finding of our study is that anterior regeneration ability has evolved several independent times among Nemertea. Mapping our regeneration dataset to nemertean phylogeny indicates four separate gains of anterior regeneration: one among Palaeonemertea and three among Pilidiophora. The origin within Palaeonemertea involves two species of the same genus (*Tubulanus sexlineatus* and *T. ruber*) that represent two fairly diverged subclades within the genus [62], indicating a gain of anterior regeneration that could be ancient. In contrast, within Pilidiophora, two gains involve a single species and the third, a pair of very closely related species, indicating that some origins of anterior regeneration within Nemertea could be relatively recent.

The number of origins of anterior regeneration in Nemertea is likely to be greater than the four formally identified in our analysis. In particular, two additional species are also reported in the literature as being capable of regenerating a full head, albeit under a narrow range of conditions: the pilidiophoran *Lineus pictifrons* [35] and the hoplonemertean *Prostoma graecense* [54]. *Lineus pictifrons* is described by Coe [35] as being able to regenerate an anterior end including the brain when amputated behind the mouth (which is posterior to the brain in this species), an observation we consider reliable given that Coe did extensive work on nemertean regeneration (including the groundbreaking work on regeneration in *L. sanguineus*). Absence of sequence information precluded us from including this species in our analysis. Because the genus *Lineus* is large, non-monophyletic [55] and includes both anteriorly regenerating and non-anteriorly regenerating species, determining whether or not *L. pictifrons* represents yet another origin of anterior regeneration must await further studies that can place this species within the nemertean phylogeny. As for *P. graecense*, this species is also reported by Kipke to regenerate a complete head [54], although only if the amputation plane is immediately behind the brain. We were unable to procure specimens of this species, precluding us from confirming this regeneration finding. However, we did have regeneration and sequence information for another species of *Prostoma*, *P. eilhardii*. Although *Prostoma eilhardii* is thought to be either very closely related to *P. graecense* or even its junior synonym [63], it showed no evidence of anterior regeneration in our experiments. Thus, if Kipke’s report is confirmed, *Prostoma graecense* would represent another very recent gain of anterior regeneration and would also indicate that gains have also occurred within the third major nemertean clade, the Hoplonemertea. If future studies corroborate these preliminary conclusions, then six gains of anterior regeneration would be inferred within Nemertea, including gains within all three major clades of the phylum.

Sampling additional nemertean species will be critical for strengthening, or revising, our understanding of the evolution of anterior regeneration in this phylum. In particular, sampling additional basal pilidiophorans is particularly important to better evaluate the ancestral condition of Pilidiophora (which in our dataset is strongly influenced by the lack of anterior regeneration in the pilidiophoran *Hubrechtia*). More extensive sampling of Paleonemertea is also needed, as only four species were included in our dataset and yet sampling within of this group is critical for confidently reconstructing the ancestral regeneration condition for Nemertea as a whole.

Finding evidence of several independent gains of head regeneration within ribbon worms suggests their body plan and biology might facilitate evolving this developmental capability. Interestingly, we documented that many nemertean species incapable of anterior regeneration can nonetheless survive without a head for an extended period of time, in some cases up to many months (Figure 2, Table S4), consistent with anecdotal observations made by other researchers. This finding is important for several reasons. First, the confidence in determining that a species fails to regenerate increases with survival time of the amputee. Second, long-term observations of amputees are crucial to assess regenerative abilities, as regeneration rates vary widely, both among and within species. And third, the ability to survive without a lost structure long enough to allow for regeneration is considered a fundamental requirement for regenerative ability to be acted upon by selection [64]. Thus, the finding that many nemertean species can survive for long periods of time without their heads certainly facilitates assessments of their anterior regeneration potential. However, and very importantly, this ability to survive without a head may be a key pre-adaptation that potentiates evolutionary gains of the ability to regenerate a head in this phylum.

### Recent gain of regenerative ability can be studied using *Lineus sanguineus* and its close relatives as a model system

The phylogenetic distribution of regenerative abilities across the Metazoa suggests that early animals, including the bilaterian stem group, were likely to have high regenerative ability [1] and that evolutionary loss of regenerative abilities appears to be far more common than gains [22]. As a consequence, our understanding of evolutionary change in regenerative ability is based almost exclusively on studies of the loss of regeneration [17,18]. Studying gains of regeneration would not only greatly improve our understanding of the developmental strategies that enable and enhance regenerative processes, but also offer insights into the organismal traits that can facilitate or constrain such gains. Unfortunately, the few previously cases of evolutionary gains of regeneration previously described map to deep branches of the Metazoan tree, and thus are too ancient to provide strong insight on the proximate causes of regeneration gains.

Of the gains identified in our study, the one represented by *Lineus sanguineus* and *L. pseudolacteus* stands out as a particularly powerful system in which to investigate the acquisition of regenerative ability. Our analysis demonstrates that the spectacular and well-documented regenerative ability of *Lineus sanguineus* emerges from a clade in which regeneration ability is relatively low, and where anterior regeneration is notably absent. The closest relatives of *L. sanguineus* are *L. lacteus*, which cannot regenerate anteriorly, and *L. pseudolacteus*, which can regenerate anteriorly although less robustly than *L. sanguineus*. Interestingly, *L. pseudolacteus*, long considered to be closely allied with *L. lacteus* and *L. sanguineus*, has recently been identified by transcriptome sequencing as a hybrid species descended by exclusive asexual reproduction from a triploid founding individual likely resulting from the fertilization of an unreduced *L. sanguineus* egg by a *L. lacteus* sperm [43]. This hybrid origin could possibly explain why *L. pseudolacteus* individuals possess regenerative abilities intermediate between those of *L. lacteus* and *L. sanguineus* [53]. Regarding the age of the gain of anterior regeneration in *L. sanguineus*, that gain is necessarily more recent than the divergence between L. sanguineus and *L. lacteus*. While no molecular clock calibration is available for nemerteans, rough estimates based on either vertebrate or protostome substitution rates suggest that this gain of anterior regeneration in the *L. sanguineus* lineage occurred within the last 10 million years [43,65].

In summary, the trio of *Lineus* species including *L. sanguineus*, *L. lacteus*, and their hybrid species *L. pseudolacteus* constitutes a powerful group in which to study the gain of regenerative ability. This species group provides an unparalleled set of advantages for future study of the evolution of regeneration: the two non-hybrid species, *L. sanguineus* and *L. lacteus*, straddle a clear gain of regeneration; the age of the regeneration gain is recent (estimated at less than 10 my); three degrees of regenerative ability is represented by the group, from non-anteriorly regenerating (in *L. lacteus*), to anteriorly regenerating in limited contexts (in *L. pseudolacteus*), to extremely robust anterior regeneration (in *L. sanguineus*); the three species are accessible, being found in similar inter- and subtidal substrates along the European coasts in reasonably large numbers to make their study convenient; and many aspects of their biology have been well described [26,37–40,57,58,66–73], providing a solid foundation on which to base new studies, including ones using the newest molecular tools.

### Systematic surveys with dense taxonomic sampling are crucial to understand how regeneration evolves

Understanding the proximate and ultimate causes of trait evolution requires confidently locating where transitions in the state of a trait have occurred across the phylogenetic history of a group. This approach involves two primary efforts: (i) documenting the state of a trait across a group of species, sampling as densely as possible; and (ii) establishing well supported hypotheses for the phylogenetic relationships among those species. For certain traits and taxonomic groups, enough information is published and available to achieve these two aims through data synthesis alone (as recently done for the study of regeneration in the phylum Annelida [23]). For other traits and taxonomic groups, the data available are too sparse to make conclusions through data synthesis alone, even if there is a long history of research on this topic [12] (as was the case for regeneration in the phylum Nemertea). In such cases, targeted data collection efforts are necessary, involving systematic surveys of the trait, taxonomic identification of specimens, and molecular phylogenetic analysis. Broad-scale efforts to reconstruct trait evolution across a large group will benefit from making use of existing, or newly established, collaboration networks and should identify mechanisms to ensure high quality of data and project integration, such as standardization of procedures and molecular barcoding. Although such efforts may seem daunting, especially when the focal group is large, they can provide important new perspectives on trait evolution, as demonstrated by this study.

Regarding the evolution of regeneration, our study clearly demonstrates that different phyla may tell different stories. Phylum-level reconstructions of regenerative ability are now available for two phyla, Annelida and Nemertea, and even though these are relatively closely related and share many aspects of morphology and natural history (being spiralian phyla composed of largely aquatic, soft-bodied, worm-like animals), these two phyla show marked differences in their evolutionary patterns of regeneration ability. These findings demonstrate the high evolvability of regenerative abilities in metaozoans. Available data thus highlight the need perform such studies in additional groups and provide strong justification for comparative studies of the developmental mechanisms underlying the evolution of regeneration.

## Data accessibility

New sequence data is being submitted to NCBI; see Table S5 in the electronic supplementary materials for accessions. Other datasets supporting the conclusions of this article will be available from the Dryad repository.

## Author’s contributions

E.E.Z., A.E.B. and J.L.N. designed the experiments and secured funding; E.E.Z., F.A.A., T.C.H. and J.L.N. collected specimens and performed regeneration experiments; E.E.Z. performed DNA sequencing experiments and analyzed the data; E.E.Z. and A.E.B. wrote the manuscript. All authors read, contributed comments on, and ultimately approved the manuscript.

## Acknowledgments

We would like to thank Svetlana Maslakova, Christopher Laumer, Susan Hill, Andrew Matthweson, Pia Floria and Carlos Vives for assisting with specimen collection, and Francesca Leasi, Herman Wirshing and the staff at the Laboratories of Analytical Biology (Smithsonian Institution, National Museum of Natural History), where DNA extraction and sequencing took place, and Jaime González Cueto for first pointing out potential regeneration in *Baseodiscus delineatus*. Some sequences were derived from a developmental transcriptome of *Prosadenoporus californiensis* sequenced and assembled by Laurel Hiebert, with support from the NSF grant IOS-1120537 to Svetlana Maslakova.

